# Copy number variation in the genome of four South American camelid species

**DOI:** 10.1101/2025.08.28.672513

**Authors:** Carola Melo Rojas, Diana Sánchez Herencia, Eloy Condori Chuchi, Japhet Zapana Pineda, Antonia Ttitto Ttica, Luis Hachircana Yauri, Víctor López Durand, Halley Rodriguez Huanca, Maximo Melo Anccasi

**Author notes:** Corresponding autor. These authors contributed equally to this work.

## Abstract

Copy number variation (CNV), a type of structural genomic variant, has been widely studied in humans, plants, and livestock. CNVs may contribute to phenotypic variation and traits of economic importance. Evaluating CNVs in domestic populations and their wild ancestors can reveal genetic changes associated with phenotypic differences that have emerged during domestication. In this study, whole-genome data from sixteen alpacas, six llamas, one guanaco, and one vicuña were used to investigate CNVs. A total of 4,247 CNVs were identified through genome-wide analysis. Functional analysis of Gene Ontology (GO) groups indicated that CNVs in alpacas are mainly related to immune-related genes, olfactory receptor genes, and keratinization. Species-level differences were observed, with the vicuña showing the least variation and the alpaca the most. This study provides the first CNV map for camelids and contributes to expanding knowledge of CNV diversity across the four South American camelid species.

## Introduction

Copy number variation (CNV) is defined as the large-scale gain or loss of DNA fragments, ranging from 50 base pairs (bp) to several megabases (Mb), relative to a reference genome. CNVs are one of the major classes of genetic variation (Zhang et al., 2009; Alkan et al., 2011). They can influence gene expression and function through multiple mechanisms, including altered gene dosage, disruption of coding sequences, and perturbation of long-range gene regulation (Qian et al., 2023). Such changes can modify gene structure and function, thereby affecting phenotypes (Weischenfeldt et al., 2013). Compared to single nucleotide polymorphisms (SNPs), CNVs exert more direct effects on gene dosage and, consequently, indirect effects on gene expression (Marques-Bonet et al., 2009).

Numerous studies have examined the relationship between CNVs and economically important traits in domestic animals. For example, CNVs affecting coat color have been identified in sheep (Norris & Whan, 2008; Fontanesi et al., 2010), goats (Fontanesi et al., 2009), cattle (Durkin et al., 2012), and pigs (Pielberg et al., 2002). CNVs have also been linked to fertility and production traits in cattle (Yue et al., 2014; Ahmad et al., 2022) and reproductive traits in pigs (Zheng et al., 2020; Qian et al., 2023; Zhang et al., 2024).

Various methodologies are available for CNV detection, including array-based comparative genomic hybridization (aCGH), SNP arrays, and whole-genome sequencing (Zheng et al., 2020; Hu et al., 2020; Liu et al., 2024). The latter is particularly useful for identifying novel and rare CNVs, especially in species for which array technologies are limited or unavailable (Jenkins et al., 2016).

South American camelids include wild species (vicuña and guanaco) and domestic species (alpaca and llama). These animals are adapted to extreme environmental conditions, including cold, high-altitude Andean regions with limited forage and low oxygen availability. The guanaco has also adapted to arid desert habitats. Alpacas and llamas are raised for meat, and alpacas additionally produce fine fiber available in a wide range of shades. Wild camelids are renowned for producing some of the finest natural fibers, a trait shaped by natural selection rather than artificial breeding (Villarreal, 2003). Despite the economic and cultural importance of camelids and recent advances in assembling the complete alpaca genome, CNV studies based on whole-genome sequencing remain scarce for this group.

The objectives of this study were to: (1) characterize CNV variation in South American camelid species, (2) identify differences between domestic and wild camelids based on whole-genome data, and (3) associate CNVs with genes and traits linked to productivity, adaptation, domestication, and other relevant phenotypes.

## Materials and Methods

Twenty-four whole-blood samples were collected from both domestic and wild camelids. Animals were selected based on phenotypic characteristics, including fiber fineness or solid coat color in alpacas and body size in llamas (Nuñoa), as well as animals with traits considered undesirable according to breed standards (CICAS La Raya, UNSAAC). Blood was collected by venipuncture into EDTA vacutainer tubes and stored at -20 °C until processing. Genomic DNA was extracted using the PureLink Genomic Mini Kit (Qiagen) according to the manufacturer’s instructions. DNA integrity and concentration were assessed using a Qubit® 3.0 fluorometer (Invitrogen, USA) and a NanoDrop spectrophotometer (Thermo Fisher Scientific, USA).

To detect CNVs, genomic DNA libraries were prepared for each sample with an insert size of ∼300 bp and sequenced on a BGISEQ-500 platform to a coverage depth of 50x. Reads were mapped to the VicPac3.1 (GCF_000164845.3) alpaca reference genome using the Burrows–Wheeler Aligner MEM algorithm. The resulting alignments were sorted by coordinate using SAMtools version 1.13 (Li et al., 2009). CNVs were detected using Control-FREEC (Boeva et al., 2012; https://boevalab.inf.ethz.ch/FREEC/). Functional annotations were obtained using Annovar (Wang et al., 2010), which classified genomic regions as exonic, intronic, upstream, downstream, or intergenic.

Gene Ontology (GO) and Kyoto Encyclopedia of Genes and Genomes (KEGG) analyses were performed using DAVID (https://david.ncifcrf.gov) (Sherman et al., 2022), and SRplot (https://www.bioinformatics.com.cn/plot_basic_pathway_enrichment_categorical_bar_plot_124_en) (Tang et al., 2023) to annotate selected genes.

## Results

A total of 4,247 copy number variants (CNVs) were detected across the 24 samples, comprising 2,622 deletions and 1,265 duplications, with an average of 175 CNVs per animal (Table 1). Data on CNVs for specific genomic regions are provided in Supplementary Table 1. CNVs were distributed across 16 of the 37 camelid chromosomes (Figure 1). Chromosome 2 contained the highest number of CNVs (407), while chromosome 13 had the fewest (109). Several chromosomes showed no CNVs in any of the analyzed samples.

**Table 1:** Copy number variants (CNVs) detected in 24 South American camelids.

| Sample | Breed | Number of deletions | Number of Duplications | Total deletion length (bp) | Total duplication length (bp) |
| --- | --- | --- | --- | --- | --- |
| White Alpaca | Huacaya | 86 | 61 | 5573328 | 4423775 |
| Black Alpaca | Huacaya | 116 | 70 | 7651037 | 4985590 |
| Black Alpaca | Huacaya | 104 | 73 | 5998673 | 4938644 |
| Brown Alpaca | Huacaya | 94 | 61 | 5794778 | 3840359 |
| Brown Alpaca | Huacaya | 86 | 66 | 5904297 | 4905061 |
| Brown Alpaca | Huacaya | 111 | 62 | 7673640 | 4567966 |
| Brown Alpaca | Huacaya | 131 | 65 | 8508857 | 4630355 |
| Spotted Alpaca | Huacaya | 126 | 62 | 7406267 | 4640358 |
| White Alpaca | Suri | 99 | 74 | 7181325 | 4342255 |
| White Alpaca | Suri | 114 | 59 | 7196279 | 3565601 |
| White Alpaca | Suri | 119 | 79 | 7637467 | 4655354 |
| Black Alpaca | Suri | 115 | 66 | 6846278 | 3835594 |
| Brown Alpaca | Suri | 105 | 62 | 7245055 | 4085598 |
| Brown Alpaca | Suri | 116 | 69 | 6596277 | 4160351 |
| LF Alpaca | Suri | 109 | 62 | 7286284 | 3667966 |
| Brown Llama | K'ara | 116 | 73 | 7606277 | 4562382 |
| Brown Llama | K'ara | 127 | 66 | 8288168 | 4265594 |
| Brown Llama | K'ara | 88 | 64 | 5463877 | 4543168 |
| Brown Llama | K'ara | 111 | 73 | 7243836 | 4515587 |
| Brown Llama | K'ara | 110 | 66 | 7438107 | 4958047 |
| Multicolor Llama | Chak'u | 111 | 66 | 7773836 | 3691384 |
| White Llama | Chak'u | 114 | 77 | 7356362 | 4750964 |
| Guanaco | — | 121 | 76 | 7656350 | 4680422 |
| Vicuña | — | 93 | 73 | 5180719 | 4212800 |

**Figure 1.**
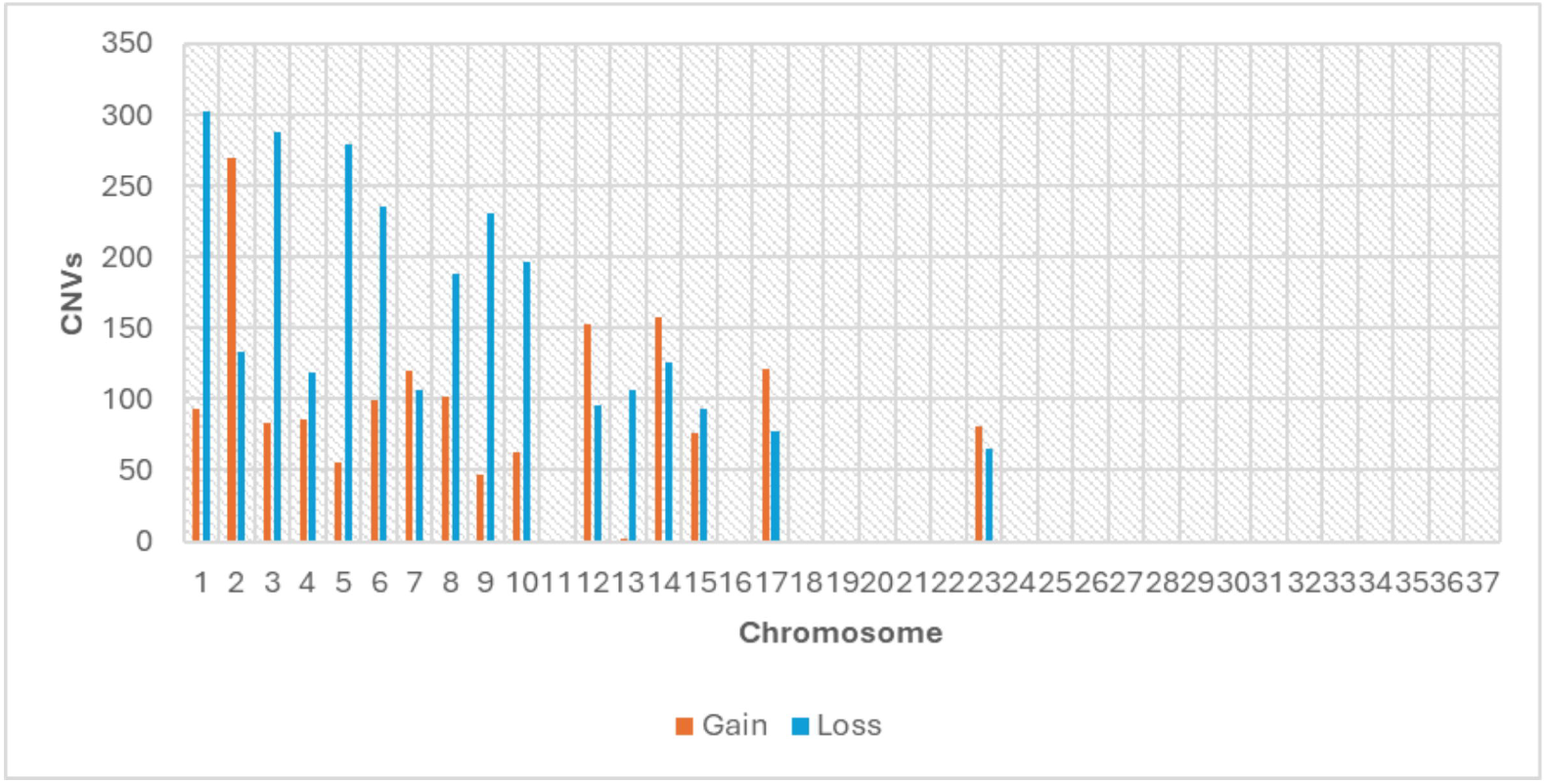

### Enrichment Analysis

To assess the functional impact of CNVs in domestic and wild camelids, Gene Ontology (GO) enrichment analysis was performed for each species using the DAVID database (Figures 2-5). Alpacas had the largest number of enriched functional groups (83), including 38 related to biological processes, 17 to cellular components, and 29 to molecular functions (Figure 2). Llamas followed with 54 enriched functional groups (26 biological processes, 11 cellular components, and 17 molecular functions) (Figure 3). Guanacos had 26 enriched functional groups (12 biological processes, 5 cellular components, and 9 molecular functions) (Figure 4), and vicuñas had 24 enriched groups (9 biological processes, 4 cellular components, and 11 molecular functions) (Figure 5).

**Figure 2.**
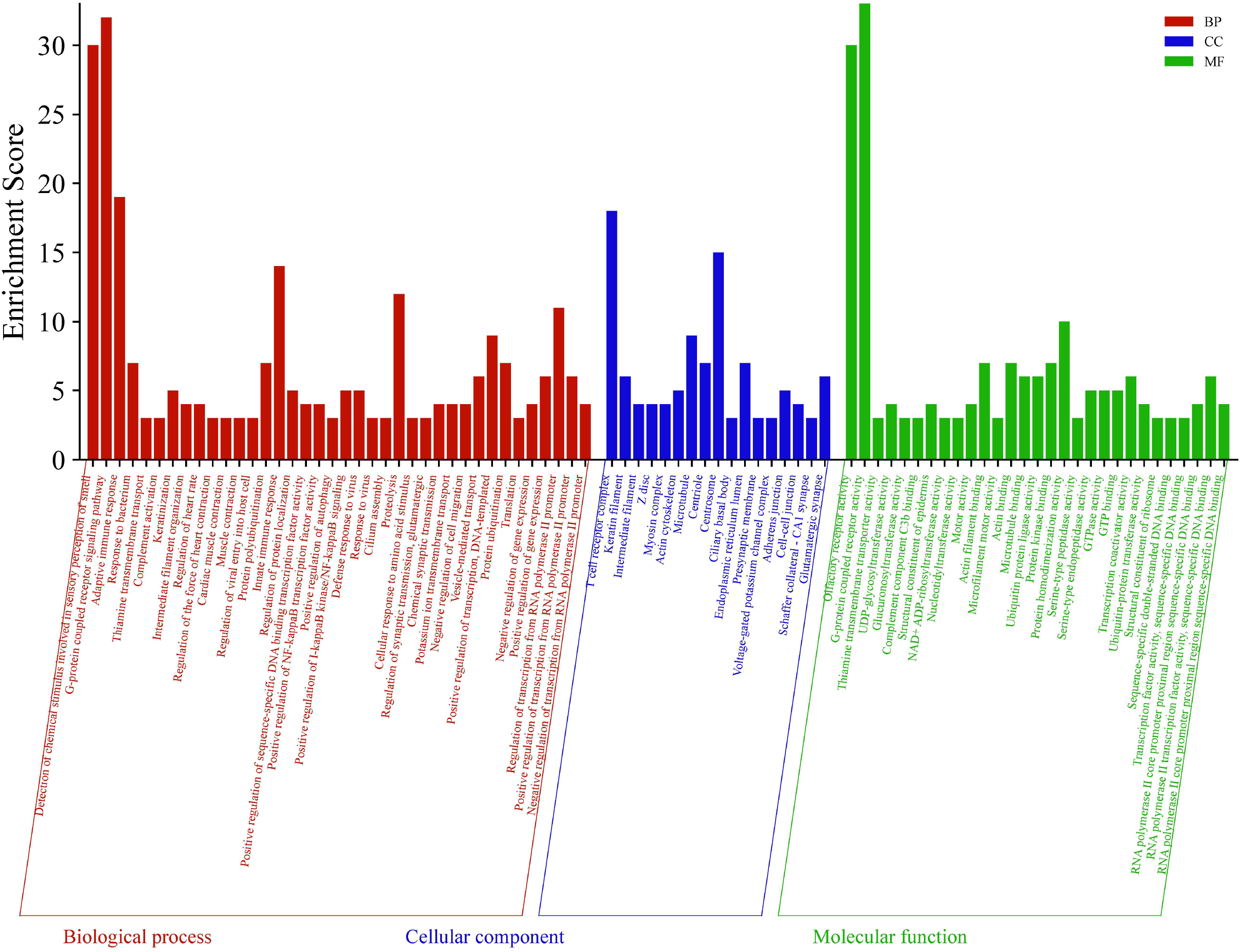

**Figure 3.**
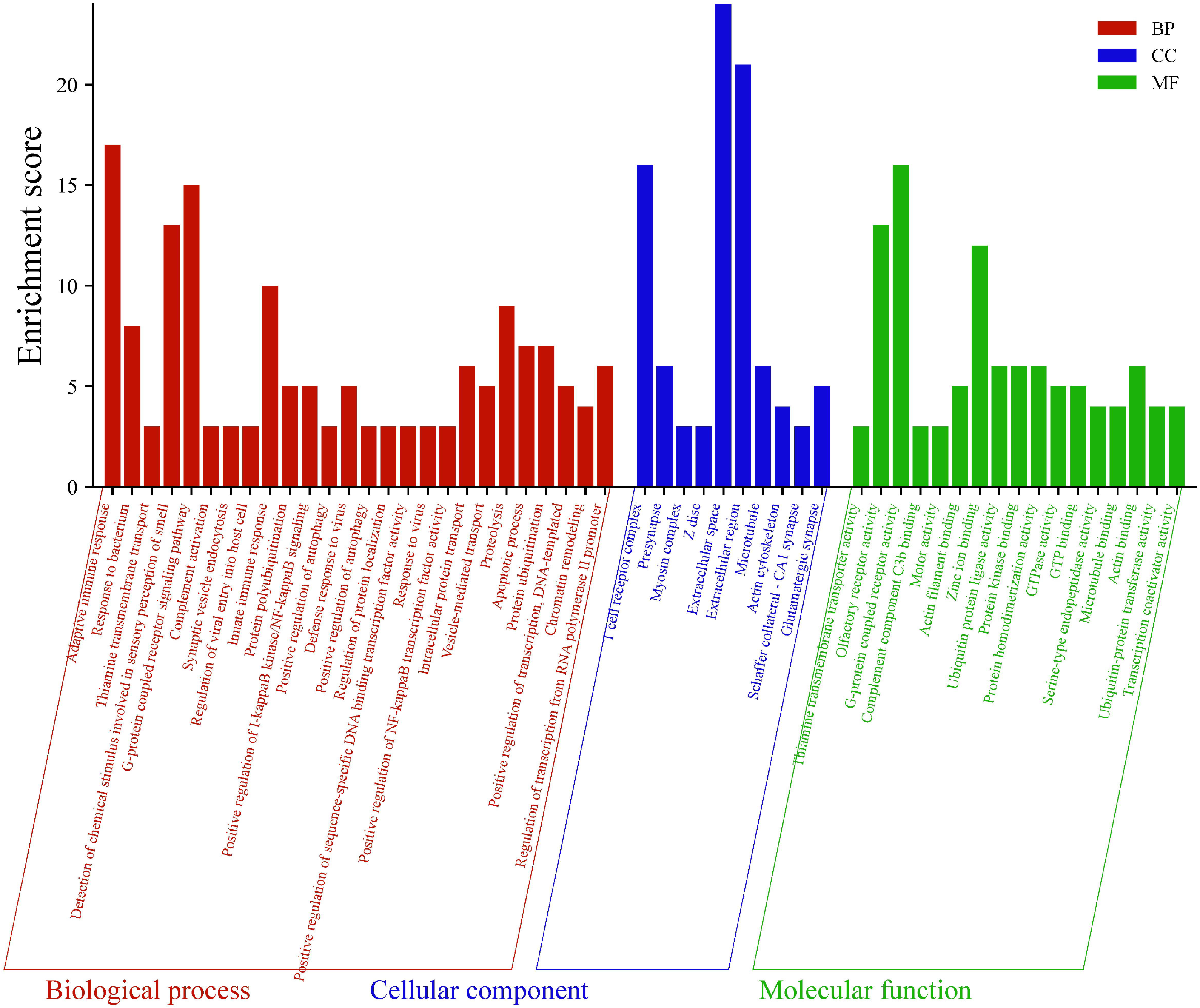

**Figure 4.**
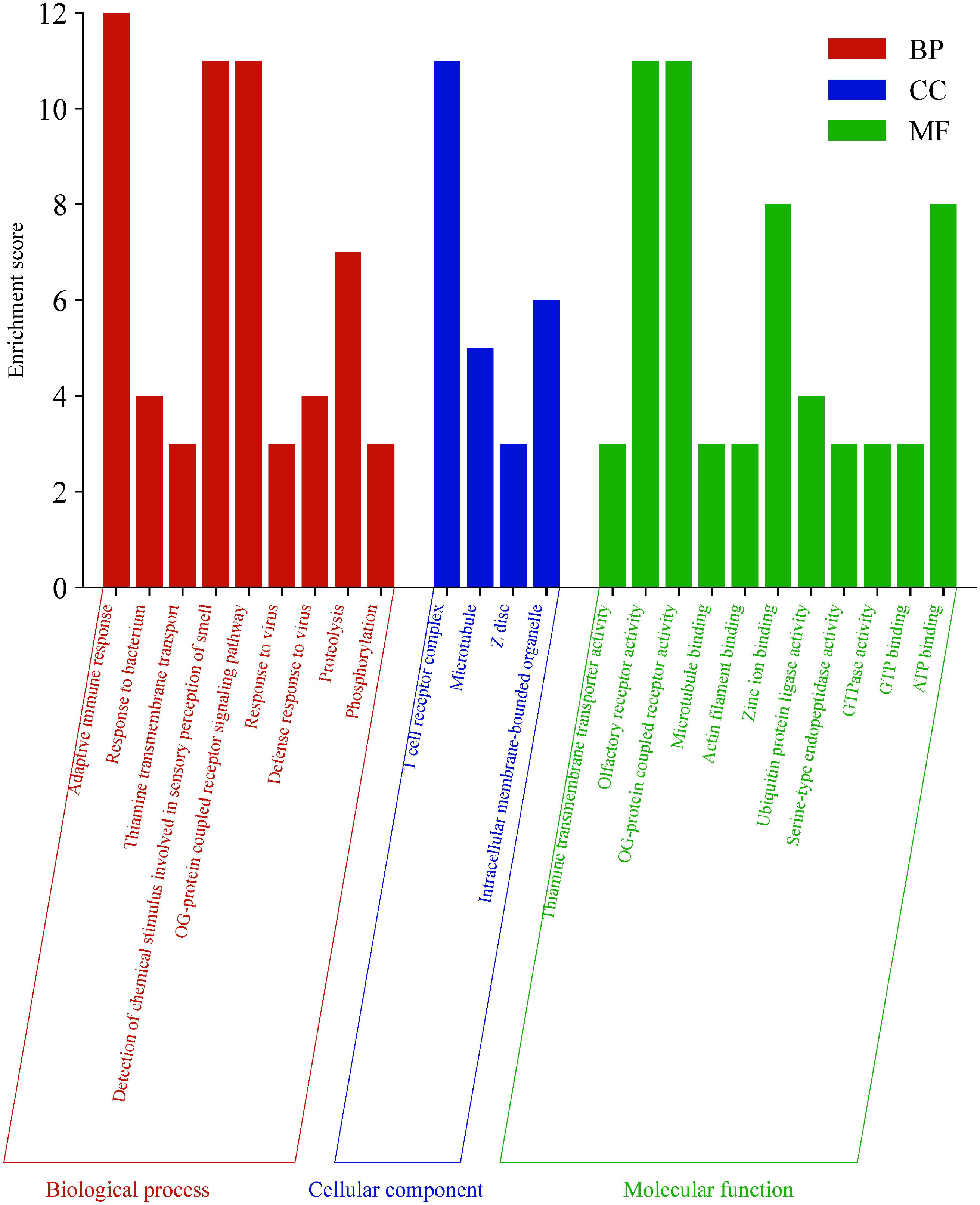

**Figure 5.**
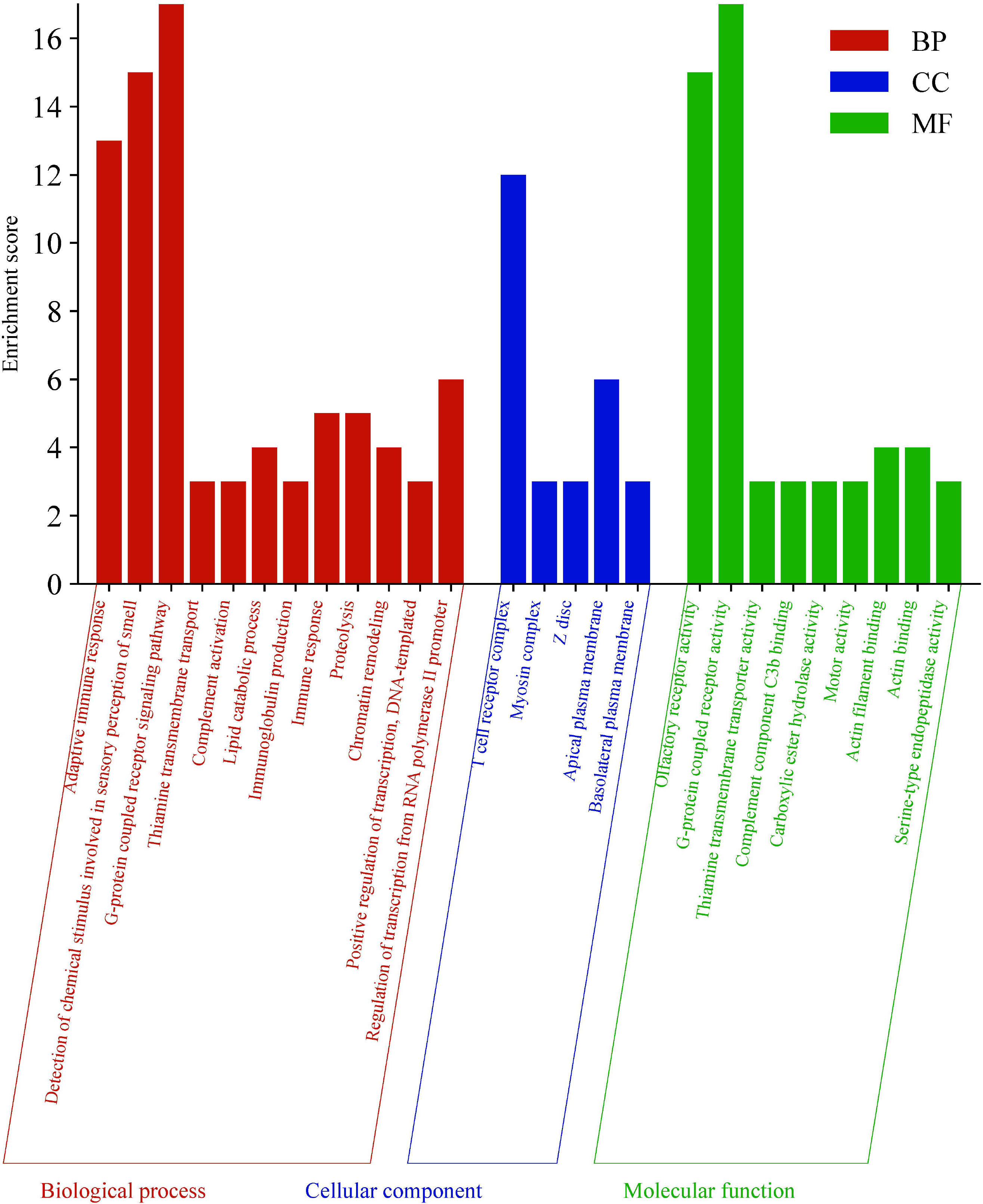

The enriched terms were largely related to adaptive immune response in all four species, innate immune response in alpacas and llamas, viral response in alpacas, llamas, and vicuñas, keratinization in alpacas, and immunoglobulin production in guanacos. These functional groups were the most prominent in the analysis. KEGG analyses identified 21 enriched pathways (Figure 6), including pathways associated with cancer, coronavirus disease, and other biological processes.

**Figure 6.**
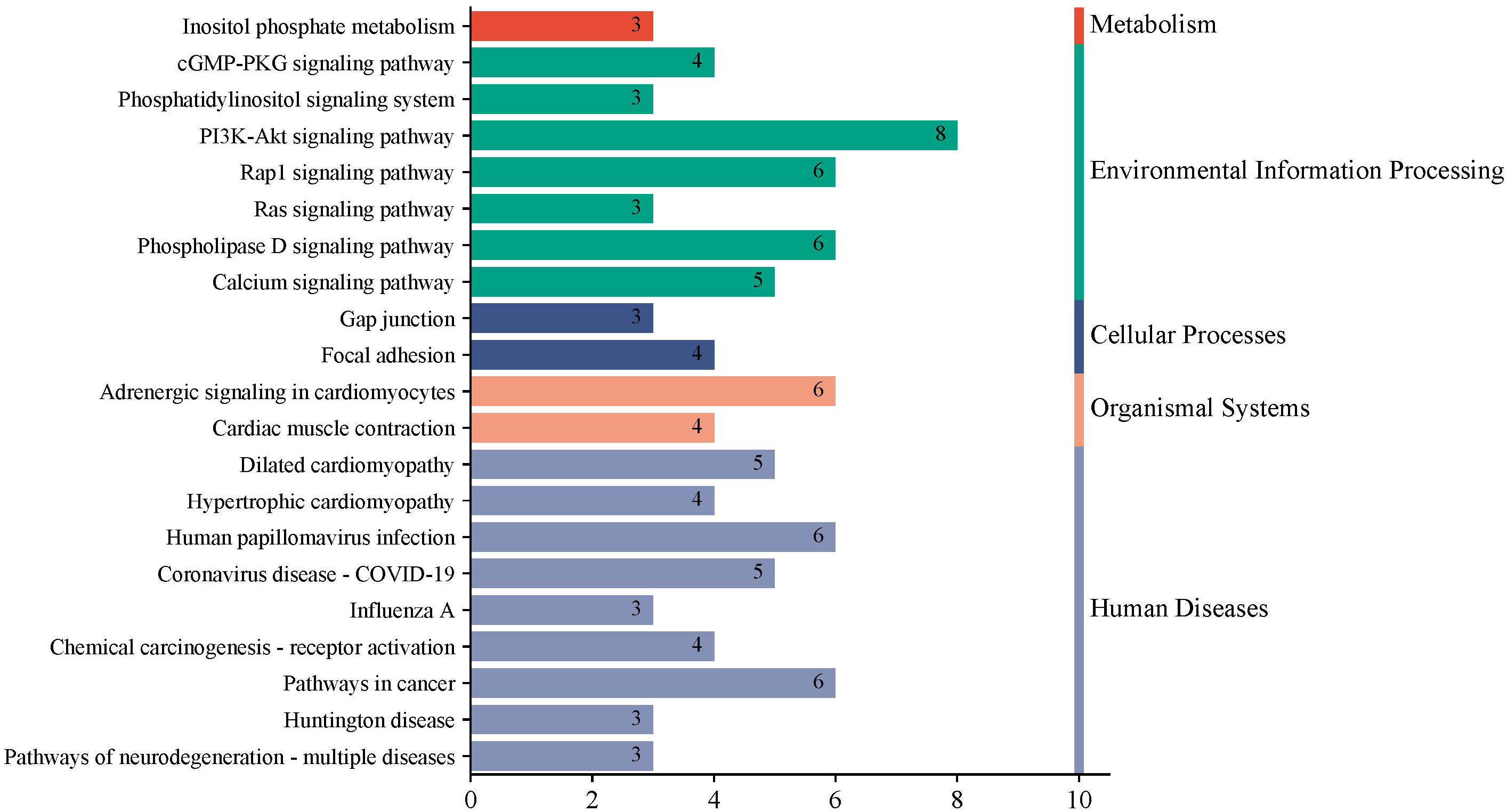

## Discussion

This study represents the first CNV analysis based on whole-genome sequencing for four South American camelid species, revealing species-specific differences in CNV profiles. On average, the CNVs identified represented 0.26% of each animal’s genome, based on alignment to the alpaca reference genome (VicPac4). This proportion is slightly lower than values reported for Tibetan sheep (0.3%; Hu et al., 2022) and six goat breeds (0.3%; Guo et al., 2018) using similar whole-genome sequencing approaches. Deletions were more common than duplications, consistent with patterns reported in other species (Upadhyay et al., 2017; Yuan et al., 2021; Qian et al., 2023).

GO enrichment analyses indicated a larger number of enriched genes in alpacas compared to other camelid species. This difference partly reflects the larger sample size for alpaca breeds (Huacaya and Suri) compared to llamas (K’ara and Chak’u) and the single sample available for vicuñas and guanacos. The small sample size for vicuñas and guanacos is due to their protected status in Peru, which restricts sampling of wild camelids.

Shared CNVs among the four species were enriched in functional categories such as sensory perception, adaptive immune response, and thiamine transmembrane transport. Similar enrichment of sensory perception genes has been reported in native Chinese fine-wool sheep (Yuan et al., 2021), Tibetan sheep (Hu et al., 2022), and yaks (Wang et al., 2019). South American camelids inhabit harsh environments with limited forage availability, particularly during the critical months of June to November, and rely heavily on sensory perception for efficient foraging. Tibetan sheep and yaks live in comparable alpine environments, which may explain these genetic similarities.

A key difference in domestic camelids was the enrichment of keratinization-related GO terms in alpacas. This finding may inform species-specific breeding strategies, as certain CNVs are subject to selection during domestication and may contribute to unique phenotypic traits (Paudel et al., 2013). In alpacas, it would be used for producing quality fibre.

The KEGG pathway analysis indicated shared CNVs associated with traits related to fiber quality, human diseases, immune function, and reproduction.

Keratin-associated proteins (KRTAPs), major structural components of the hair shaft, are specifically expressed in hair shaft layers (Rogers et al., 2006). This family of genes shows considerable diversity and plays a critical role in determining fiber characteristics. The KRTAP13-1 gene, located on chromosome 1 (Mendoza et al., 2021), has been identified as a candidate gene for fiber growth in alpacas (Flórez, 2016) and has been associated with reduced wool diameter in sheep (Wua et al., 2022). Studies of the KRTAP10-1 gene in humans revealed expression in the keratinizing zone above the hair shaft cuticle (Fujikawa et al., 2013), while variants of the KRTAP20-2 gene have been linked to fiber weight in goats (Wang et al., 2017) and crimping in sheep (Mohamadipoor et al., 2021). Keratins (KRTs) are essential for forming the intermediate filament cytoskeleton in epithelial cells (Li et al., 2024). In goats, differences in KRT5 and KRT83 alleles and genotypes are associated with coat curliness (Duan et al., 2022), and differential KRT5 expression has been observed between primary and secondary follicles (Li et al., 2024). In sheep, an allele of KRT86 is associated with improved fiber structure (very fine bundles), resulting in desirable wool traits such as well-defined crimp and low variation along the fiber. These traits are relevant to wool manufacturing and have been linked to other genes, including WNT1, MPC, and RSC1A1 (Bolormaa et al., 2017).

Identifying genes associated with disease resistance or susceptibility, especially those of zoonotic importance, is crucial for breeding strategies. For instance, the MX1, MX2, and TMPRSS2 genes have been studied in patients with severe SARS-CoV-2 infections, with MX1 showing protective effects (Bizzotto et al., 2020). The 60S ribosomal protein L29 (RPL29) gene exhibits high expression in patients with SARS-CoV-2 infection and has been proposed as a potential nanomedicine target (Khalid et al., 2022). A robust immune system is essential for effective disease defense, particularly in wild camelids. Lysozyme (LYZ), an antimicrobial enzyme, is a key component of the innate immune system (Glamočlija et al., 2020); in buffalo, its expression in the mammary gland is significantly associated with antibacterial activity (Su et al., 2023). In pigs, copy number variation in LYZ is associated with immune system traits (Zhang et al., 2024).

This study also identified a CNV in the CATSPERε gene, which is essential for sperm hyperactivation and male fertility. CATSPERε-deficient males are sterile due to the absence of a complete CATSPER channel and defective sperm motility (Hwang et al., 2025). The CNV involving this gene was observed only in the vicuña sample, and further research is required to clarify its role in reproductive

## Conclusion

In this study, genome sequencing data from four South American camelid species were used to detect copy number variations (CNVs), enabling comparisons between domestic and wild populations. Several CNVs were identified in genes associated with fiber characteristics (KRTAP10-1, KRTAP13-1, KRTAP20-2, KRT5, KRT83, KRT86), immune function (MX1, MX2, RPL29, LYZ), and reproduction (CATSPERε). These genes may influence economically important traits such as fiber quality in alpacas and llamas, immune system performance, and reproductive capacity.

The results of this study provide a valuable resource for understanding genomic variation in alpacas, llamas, guanacos, and vicuñas. They also offer a foundation for future research on trait-associated CNVs, and selective breeding strategies aimed at improving productivity and adaptation in domestic camelids.

## Supporting information

Supplementary Tdocuments

