## Supplementary Tdocuments for "Copy number variation in the genome of four South American camelid species": Tabla_sup_1.docx

| Number | Sample | Breed | Upstream | Exonic | Intronic | Splicing | Downstream | Upstream/Downstream | UTR5 | UTR3 | Intergenic |
| --- | --- | --- | --- | --- | --- | --- | --- | --- | --- | --- | --- |
| 1 | White Alpaca | Huacaya | 2 | 85 | 6 | 0 | 0 | 0 | 2 | 1 | 50 |
| 2 | Brown Alpaca | Huacaya | 2 | 87 | 8 | 0 | 0 | 0 | 2 | 1 | 72 |
| 3 | Brown Alpaca | Huacaya | 2 | 94 | 11 | 0 | 0 | 0 | 2 | 1 | 85 |
| 4 | Black Alpaca | Huacaya | 2 | 101 | 6 | 0 | 0 | 0 | 1 | 1 | 73 |
| 5 | Black Alpaca | Huacaya | 2 | 87 | 8 | 0 | 0 | 0 | 2 | 1 | 75 |
| 6 | Brown Alpaca | Huacaya | 2 | 77 | 5 | 0 | 0 | 0 | 2 | 1 | 67 |
| 7 | Brown Alpaca | Huacaya | 1 | 78 | 6 | 0 | 0 | 0 | 2 | 1 | 63 |
| 8 | Spotted Alpaca | Huacaya | 2 | 89 | 6 | 0 | 0 | 0 | 4 | 1 | 83 |
| 9 | White Alpaca | Suri | 2 | 87 | 7 | 0 | 0 | 0 | 2 | 1 | 71 |
| 10 | Black Alpaca | Suri | 2 | 85 | 6 | 0 | 0 | 0 | 2 | 1 | 83 |
| 11 | Brown Alpaca | Suri | 2 | 85 | 7 | 0 | 0 | 0 | 2 | 1 | 68 |
| 12 | Brown Alpaca | Suri | 2 | 84 | 11 | 0 | 0 | 0 | 2 | 1 | 83 |
| 13 | LF Alpaca | Suri | 2 | 86 | 8 | 0 | 0 | 0 | 1 | 1 | 72 |
| 14 | White Alpaca | Suri | 2 | 98 | 8 | 0 | 0 | 0 | 4 | 1 | 84 |
| 15 | White Alpaca | Suri | 2 | 87 | 9 | 0 | 1 | 0 | 2 | 1 | 69 |
| 16 | Brown Llama | K'ara | 2 | 94 | 9 | 0 | 0 | 0 | 2 | 1 | 79 |
| 17 | Brown Llama | K'ara | 2 | 95 | 8 | 0 | 0 | 0 | 2 | 1 | 82 |
| 18 | Brown Llama | K'ara | 2 | 83 | 6 | 0 | 0 | 0 | 2 | 1 | 57 |
| 19 | Brown Llama | K'ara | 2 | 101 | 8 | 0 | 0 | 0 | 2 | 1 | 69 |
| 20 | Brown Llama | K'ara | 2 | 93 | 7 | 0 | 0 | 0 | 2 | 1 | 69 |
| 21 | Multicolor Llama | Chak'u | 2 | 88 | 9 | 0 | 0 | 0 | 2 | 1 | 72 |
| 22 | White Llama | Chak'u | 2 | 96 | 9 | 0 | 0 | 0 | 1 | 1 | 80 |
| 23 | Guanaco |  | 2 | 100 | 9 | 0 | 0 | 0 | 4 | 1 | 79 |
| 24 | Vicuña |  | 2 | 95 | 4 | 0 | 0 | 0 | 3 | 1 | 59 |
